# Metabarcoding and microtomography reveal new insights into the dietary niche of near-extinct amphibians

**DOI:** 10.1101/2025.11.10.687692

**Authors:** Amadeus Plewnia, Renaud Boistel, Christopher Heine, Tobias Hildwein, Amanda B. Quezada-Riera, Andrea Terán-Valdez, Juan P. Reyes-Puig, Stefan Lötters

## Abstract

Harlequin toads (*Atelopus*) are among the most threatened vertebrates, yet their trophic ecology remains poorly understood due to the virtual disappearance of most populations and non-invasive sampling constraints. Here, we combine DNA metabarcoding of faecal samples from six surviving harlequin toad species from Ecuador with synchrotron-based microtomography of historic, fluid-preserved material of seven species across all major clades of the genus to assess dietary composition. Metabarcoding revealed a diverse invertebrate diet with marked ecological segregation between habitats, suggesting specialization on Hymenoptera in species inhabiting low to mid elevation forests and broader prey spectra in high-Andean taxa. Synchrotron scanning for the first time allowed non-destructive recovery of 3D-images of arthropod exoskeletons from the intestinal system of amphibians, confirming hymenopterans as key prey for forest-associated *Atelopus* in historical specimens. This dual approach overcomes the limitations of traditional and single-method studies, offering a scalable, non-invasive framework for dietary analyses. By providing curated step-by-step commands for sequence processing, we further aim to make dietary metabarcoding more accessible to zoologists. Our study substantially expands dietary data for near-extinct harlequin toads and supports conservation efforts with urgently needed ecological insights as a baseline for adapted conservation breeding and integrative conservation of trophic webs.

## Introduction

As major generalist predators of invertebrates at the interplay between aquatic and terrestrial ecosystems, amphibians hold a crucial function in community webs and associated ecosystem services (Whiles et al. 2006; Colón-Gaud et al. 2009; Vignoli and Luiselli 2012; Springborn et al. 2022). At the same time, amphibians are declining at an accelerated pace globally leaving little time for researchers to document their role in ecosystem function (Stuart et al. 2004; Mendelson et al. 2006; Luedtke et al. 2023).

Harlequin toads (*Atelopus*) are a species-rich group of true toads (Bufonidae) and have radiated over much of tropical South America and adjacent Central America (e.g. Lötters 1996, Lötters et al. 2025). While the genus was once widespread across its range, in recent decades, harlequin toads have suffered rapid, unparalleled declines from the spread of emerging infectious disease and habitat loss (La Marca et al. 2005; Lötters et al. 2023). As a result, the group is among the world’s most threatened vertebrate genera today (Lötters et al. 2023; IUCN 2025). Being diurnal, colourful, and in some cases of cultural importance, harlequin toads can be considered ‘flagship’ species for Neotropical amphibian conservation and vast efforts have been undertaken to rediscover and protect the remaining populations, which ultimately has resulted in the creation of a multinational conservation network, the *Atelopus* Survival Initiative, ASI (Valencia and Marin da Fonte 2022). Conserving these emblematic amphibians heavily relies on ex situ breeding (Gratwicke et al. 2015; Lewis et al. 2019) as so far, no efficient mitigation strategies are available for the emerging infectious disease chytridiomycosis, the most severe threat to harlequin toads (Lips et al. 2008; Scheele et al. 2019; Lötters et al. 2023). The virtual absence of healthy populations in situ has hampered thorough studies of their ecology and life history including their dietary composition. However, this information is crucial for successful conservation breeding and reintroduction (cf. Lassiter et al. 2020; cf. Klocke et al. 2023). Dietary studies in *Atelopus* species number few and so far, they have relied on visual examination of arthropod body parts retained from faecal samples (e.g. Vega-Yánez et al. 2024), gastric lavage (L. A. Rueda-Solano unpubl. data) or specimen dissection (e.g. Durant and Dole 1974a, b; Toft 1981; Duellman and Lynch 1988; González et al. 2012). These methods, however, are partially invasive making them barely applicable to the few remaining populations or the small series of preserved voucher material and are further incomplete in characterizing dietary composition due to rapid degradation of soft-bodied prey.

Instead, metabarcoding of faecal and intestinal samples has recently emerged as a non-invasive, cost-efficient and standardized tool to rapidly approach and transform our understanding of community ecology and particularly to place focal species in the communities’ food web of various animal taxa (de Sousa et al. 2019; Ficetola et al. 2019; Kennedy et al. 2020).

Beyond molecular approaches, microtomography has become an emerging tool in zoological studies, particularly in systematics, functional morphology and developmental biology (Boistel et al. 2013; Clark and Badea 2021; Schubnel et al. 2022). Recent technological advances, especially phase-contrast synchrotron scanning, allow unprecedented imaging resolution of internal structures in fluid-preserved material (Sena et al. 2022). While increasingly adopted to study skeletal anatomy or soft tissue organization, microtomography has not yet been explored for dietary research in amphibians.

We here use metabarcoding of faecal samples collected from rediscovered and surviving contemporary populations to disentangle dietary niche information of harlequin toads across Amazonian and Andean Ecuador from lowland rainforest to paramo. We supplement dietary sampling gaps by employing X-ray phase contrast microtomography as a novel, non-invasive tool to visualize and identify prey remains in the digestive tract of additional historic museum specimens. This opens up a novel, non-destructive avenue for predator-prey studies in taxa where destructive methods are unfeasible or ethically restricted.

## Material and methods

### Faecal sampling

Over the course of visual and acoustic monitoring to trace lost or critically endangered populations of harlequin toads in Ecuador, we collected non-invasive faecal samples for dietary metabarcoding from six species of *Atelopus* (Fig. 1) from different habitats (paramo and sub-paramo: *A. bomolochos, A. ignescens;* cloud forest: *A. palmatus*, and, in sympatry, *A. coynei* and *A. lynchi*; lowland rainforest: *A. colomai*). All specimens encountered were captured with sterile nitrile gloves and kept in UV-sterilized plastic bags with sufficient humidity overnight. Subsequently, frogs were released at the site of capture with any faecal pellets being transferred with a sterile rayon swab and preserved in 300µl DNA/RNA Shield (Zymo Research, Freiburg) to prevent degradation of nucleic acids at ambient conditions. Species and specimens sampled are detailed in Supplementary Material. Fieldwork and sample collection was granted by Ministerio del Ambiente, Agua y Transicciones Ecologicos del Ecuador (Quito) under code No. MAATE-DBI-CM-2022-0261 and MAATE-ARSFC-2022-2344.

**Figure 1.**
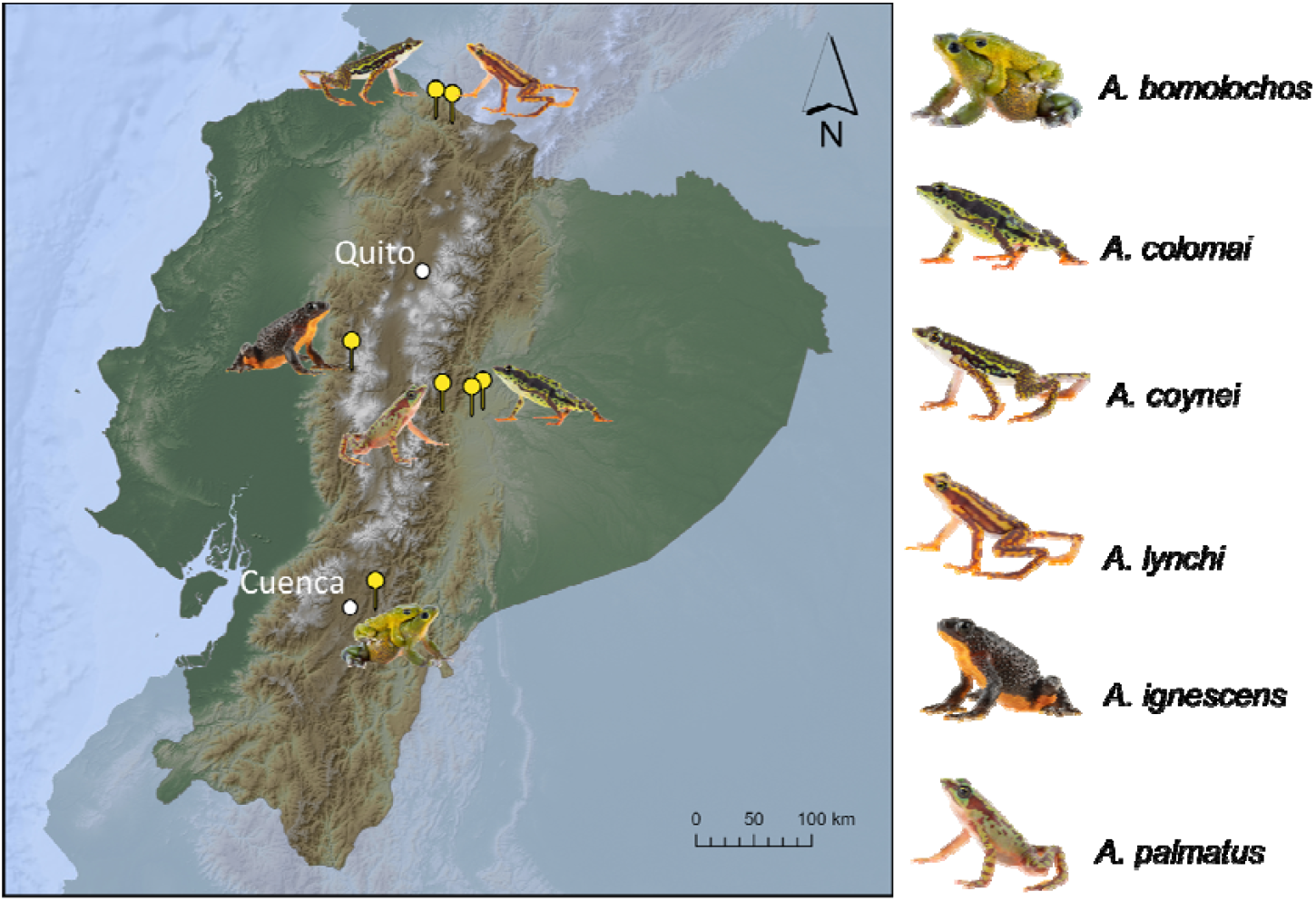
Altitudinal map of Ecuador with sampling sites of harlequin toad species studied with metabarcoding. Photos: Jaime Culebras (Photo Wildlife Tours) and Amanda B. Quezada-Riera.

### Library preparation and sequencing

Work was conducted in an UV-sterilized laminar flow hood to minimize contamination risk with field negative (rayon swab without faecal pellet preserved in the field) and lab negative (RNAse-free water in extraction and PCR) samples processed alongside all samples. We extracted genomic DNA using the QIAamp PowerFecal Pro kit (Qiagen, Hilden) following the manufacturer’s instructions. We targeted and amplified a 116 bp fragment of the mitochondrial barcoding region COI using the arthropod-specific primer pair NoPlantF_270 (5’-RGCHTTYCCHCGWATAAAYAAYATAAG-3’) and mICOIintR_W (5’-GRGGRTAWACWGTTCAWCCWGTNCC-3’) with primers carrying Illumina TruSeq adapters for subsequent indexing (Krehenwinkel et al. 2022). Amplicons were coded with dual indices (Illumina TruSeq i7 and i5, Illumina) in a subsequent PCR with 6 cycles following Lange et al. (2014). Each DNA extract was amplified in duplicates and each replicate indexed individually. Indexed PCR products were pooled in equal amounts into a single library and purified using NucleoMag magnetic beads (Macherey-Nagel, Düren) following the manufacturer’s instructions. We used the MiSeq Reagent Kit v2 with 500 cycles for sequencing on a MiSeq platform (Illumina, San Diego) aiming at ∼20,000 reads coverage for each replicate.

### Sequence analysis

Processing of demultiplexed paired-end MiSeq reads was conducted as described in Plewnia et al. (2025a) to generate 97% radius operational taxonomic units (OTUs). We mapped OTUs to a local copy of NCBI GenBank’s nt database (www.ncbi.nlm.nih.gov/Genbank; downloaded 20 December 2023) using Blastn and assigned taxonomic information using blast2taxonomy (Schöneberg 2024). We filtered OTU tables for Metazoa but excluding Chordata (host reads) and removed OTUs with reads ≤ 5 as described in Plewnia et al. (2025b).

Further, as we only targeted higher taxonomic levels, we included all BLASTn hits with ≥ 90 % sequence identity. After filtering, all OTUs remaining in negative controls were not found in any sample and were subsequently removed. We analyzed and visualized data in R version 4.2.2 using the packages vegan version 2.6-8 and ggplot2 version 3.4.2 (Wickham 2016; Oksanen et al. 2025). For non-metric multidimensional scaling (NMDS), we transformed data to presence/absence information and plotted prey on order level. In an effort to make our approach more accessible for other researchers, we provide detailed scripts for sequence processing and analysis as Supplementary Methods.

### Microtomography

We obtained three-dimensional data of prey item exoskeletons by X-ray phase contrast microtomography of eight preserved harlequin toad specimens corresponding to seven species from European museum collections (Supplementary Material). Specimens were scanned on the ANATOMIX beamline (Weitkamp et al. 2022) at Synchrotron SOLEIL with a central energy of the beam around 45 keV and 0.9 m distance to the detector. Scanning followed Langer et al. (2010) and Boistel et al. (2013). Scans were processed as described in Lötters et al. (2025) using the Paganin algorithm to retrieve the phase. Three-dimensional processing and rendering were performed in Avizo3D 2022.1 (Thermo Fisher Scientific). See Supplementary Material for a list of the material examined and detailed images of stomach content).

## Results

### Metabarcoding

After quality filtering of raw reads, we retrieved about 1,138,000 reads from our COI library with about 164,000 remaining reads corresponding to 254 OTUs after filtering blasted sequences. Sequencing showed saturation (i.e. deep read coverage of most OTUs; Supplementary Material). The proportion of arthropod reads among all reads was generally high (60.4 %) and each *Atelopus* specimen sampled yielded mean 5,115 (range: 9– 33,222) reads corresponding to on average 8.6 (range: 1–28) OTUs. Besides Arthropoda, several other metazoan phyla were present in the OTU dataset, among which Annelida (Clitellata), Mollusca (Gastropoda) and Onychophora were included for downstream analysis as they may constitute potential amphibian prey and are found in the general area (Supplementary Material).

The diet of the six *Atelopus* species studied mostly consisted of insects and to a lesser extent arachnids, while in addition, gastropods comprised a considerable portion in two paramo and cloud forest species, *A. bomolochos* and *A. lynchi*, respectively, and clitellate annelids in the paramo inhabitants *A. bomolochos* and *A. ignescens* (Fig. 2A). The ingested arthropods constituted mainly Coleoptera (Staphylinidae), Lepidoptera (Noctuidae), Araneae (Theridiidae) and Siphonaptera (Ischnopsyllidae) in *A. bomolochos*, Hymenoptera (Formicidae), Coleoptera (Cerambycidae), and Lepidoptera (Tortricidae, but only in one of two populations) in *A. colomai*, Lepidoptera (Oecophoridae) and Coleoptera (Buprestidae) in *A. ignescens*, Hymenoptera (Formicidae), Lepidoptera (Crambidae, Sesiidae), Coleoptera (several families), Araneae (Araneidae) and Mesostigmata (Phytoseiidae) in *A. lynchi* and Hymenoptera (Formicidae), Araneae (Theridiidae), Diptera (Chloropidae, Tipulidae), Trombidiformes (Hydryphantidae) and Orthoptera (Gryllidae) as well as an onychophoran in *A. palmatus* (Fig. 2B,C). The only sample of *A. coynei* sequenced contained a single taxon attributed to scaffold web spiders (Araneae: Nesticidae; Fig. 2B,C). For detailed dietary composition and higher taxonomic resolution see Supplementary Material.

**Figure 2.**
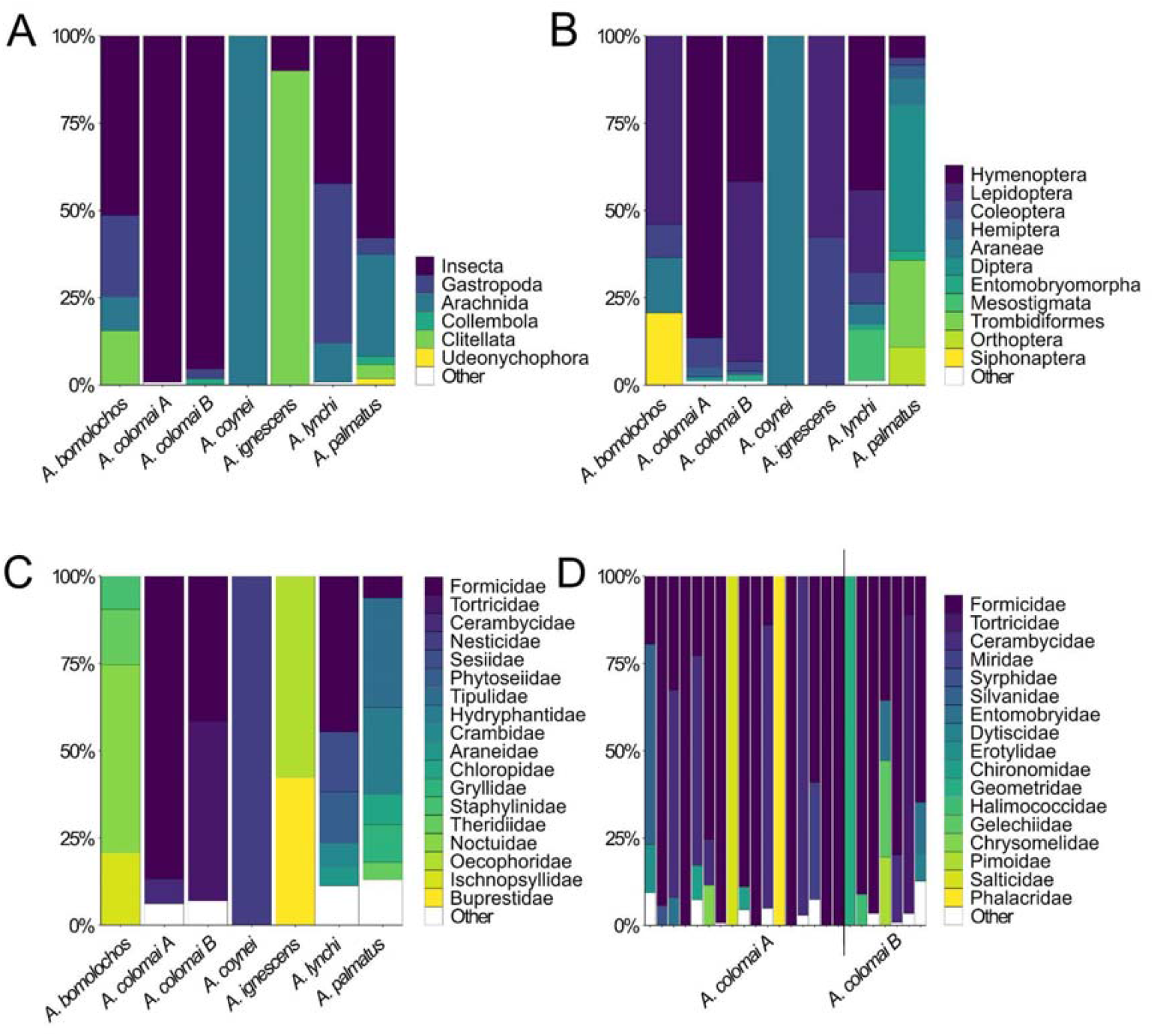
Relative read occurrence of harlequin toad prey in faecal samples. A) Metazoan groups, B) orders of Arthropoda and C) families contained in the orders displayed in B) for six species of *Atelopus* from different habitats in Ecuador. D) shows individual variation among two populations of *A. colomai*, the species best sampled in this paper.

In the case of the Amazonian lowland *A. colomai*, a more detailed sampling from two localities was available (Fig. 1), allowing to assess within- and between-population variation (Fig. 2D). In this species, while high variation in relative read abundance of families occurs within both populations, ants constituted the main prey of both populations dominating the dietary reads of 58.3 % of all individuals and being present with at least 5% of all reads in 83.3 % of all individuals (Fig. 2D).

### Microtomography

The relatively high degree of specialisation on Hymenoptera in lowland to cloud forest *Atelopus* is further corroborated by µCT imaging. The only available specimen of *A. colomai* contained a large number of ants corresponding to various morphospecies incl. undetermined Formicidae and one member of *Odontomachus*, as well as one Myriapoda, one mite and two beetles. Similarly, the stomach content visualized in the single specimen available for *A. spumarius* from the Peruvian Amazon lowlands constituted large numbers of ants and small beetles only (Fig. 3; Supplementary Material). In *A. hoogmoedi*, the single specimen investigated contained six termites (incl. Nasutitermitinae soldiers) and one degraded item likely corresponding to Formicidae (Fig. 4; Supplementary Material). In *A. tricolor* from the Amazonian Andean versant of Bolivia and Peru, the stomach contained a large number of ants with several mites (Uropodina, Phthiracaroidea), and single small specimens of Coleoptera and Hemiptera each (Supplementary Material). Both specimens of *Atelopus cruciger* from the Venezuelan Andean cloud forests similarly preyed on small Coleoptera (incl. Curculionidae and Scolytinae). In one specimen, additional Hymenoptera (incl. Isoptera, Formicinae, Ponerinae, and a likely member of Apoidea) were present as well as a small arachnid (Fig. 4). In the paramo species *A. carrikeri* from the basal Sierra Nevada clade, we found several larger Coleoptera (incl. an adult Curculionidae and a larval Elateridae), several Hymenoptera (incl. Platygastridae), one specimen corresponding to Araneae as well as a small leaf and several Nematodes (Fig. 4). In *A. ignescens* from the páramos of central Ecuador, we detected several larger Coleoptera (Carabidae and Curculionidae), Blattodea, Isoptera, a Lepidoptera larva, several Acari and an unidentified arachnid potentially belonging to Opiliones (Fig. 4).

**Figure 3.**
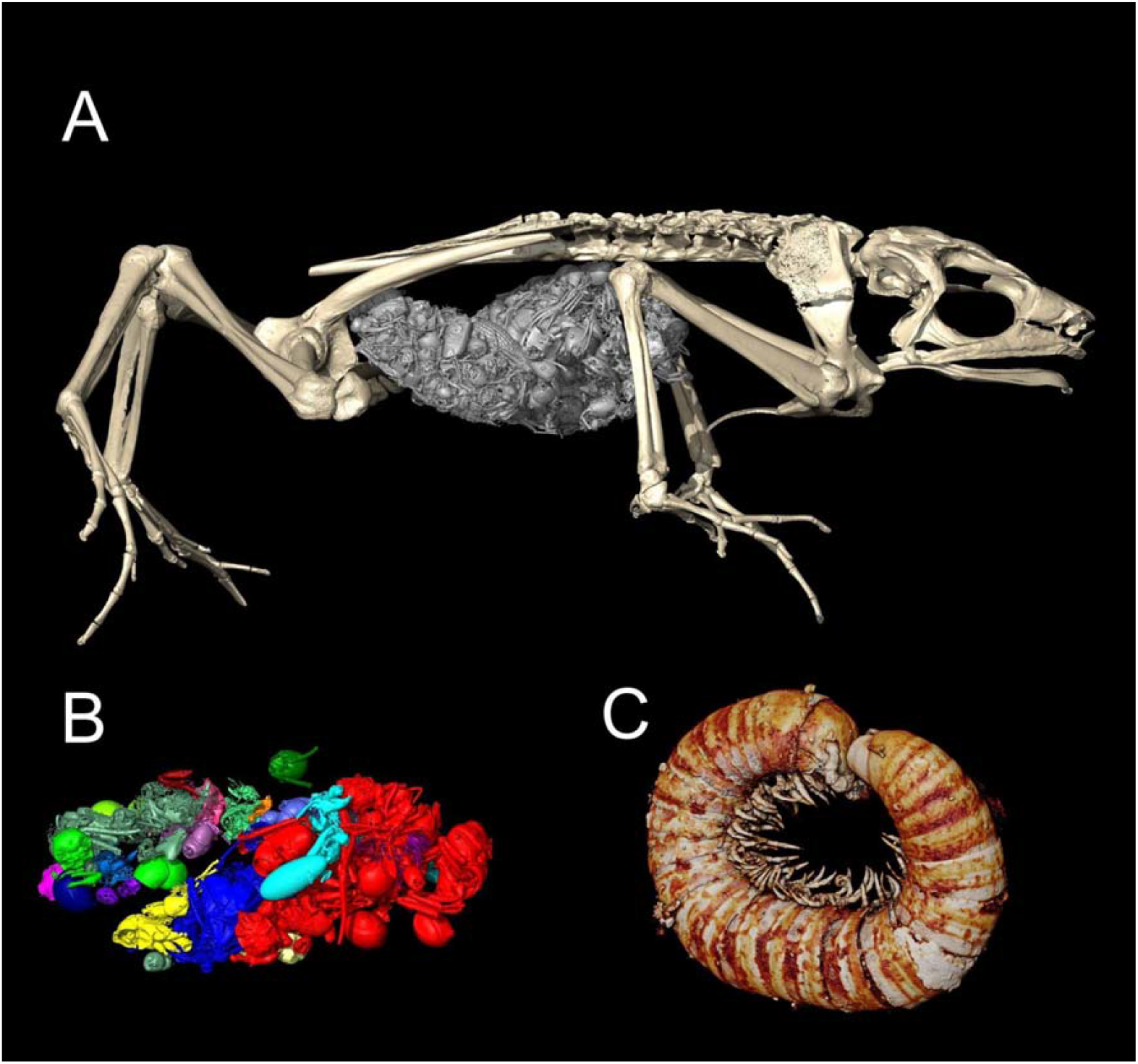
Diet of A) *Atelopus spumarius* (neotype MNHNP 1979/8382) and B, C) *A. colomai* (B with stomach content and C buccal content; paratype ZFMK 44976) visualized using synchrotron microtomography. In line with metabarcoding, synchrotron scanning recovered Formicidae as main prey with additional Coleoptera and a single Diplopoda in the case of *A. colomai*. See Fig. 4 for detailed prey items. Not to scale.

**Figure 4.**
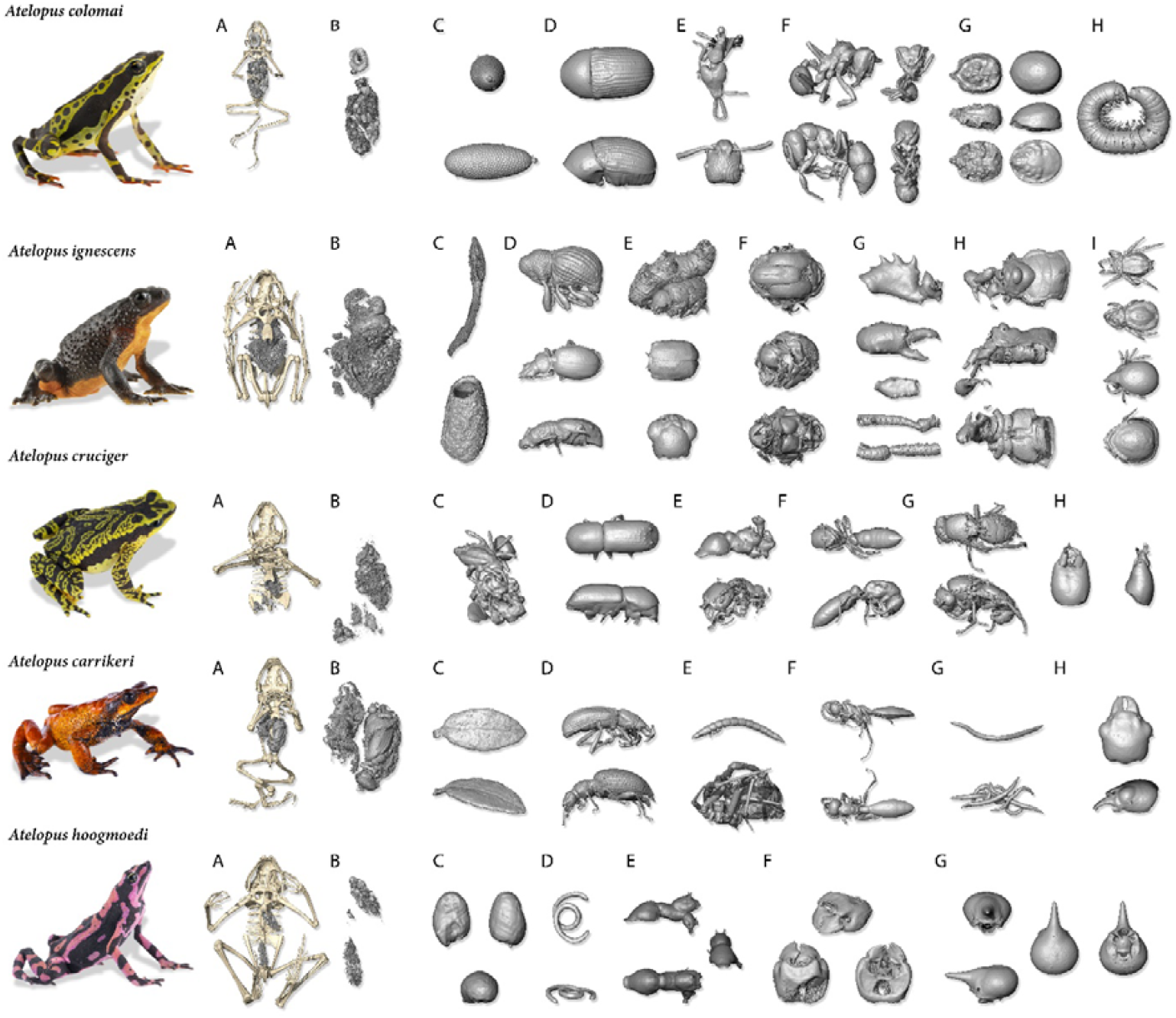
Selected individual prey items recovered with synchrotron microtomography from five *Atelopus* species across all major clades and habitat types. A) shows prey items with relative position to the amphibian specimen and B) the extracted stomach. C)-I) details single prey items extracted from imaging data. Not to scale.

Non-metric multidimensional scaling (NMDS) of metabarcoding data (Fig. 5) revealed moderate differentiation in the dietary niche of the species sampled (ANOSIM: R = 0.271, p = 0.0083) but no differentiation between populations of *A. colomai* (p > 0.05). However, the small sample size prohibits pairwise comparisons for most species.

**Figure 5.**
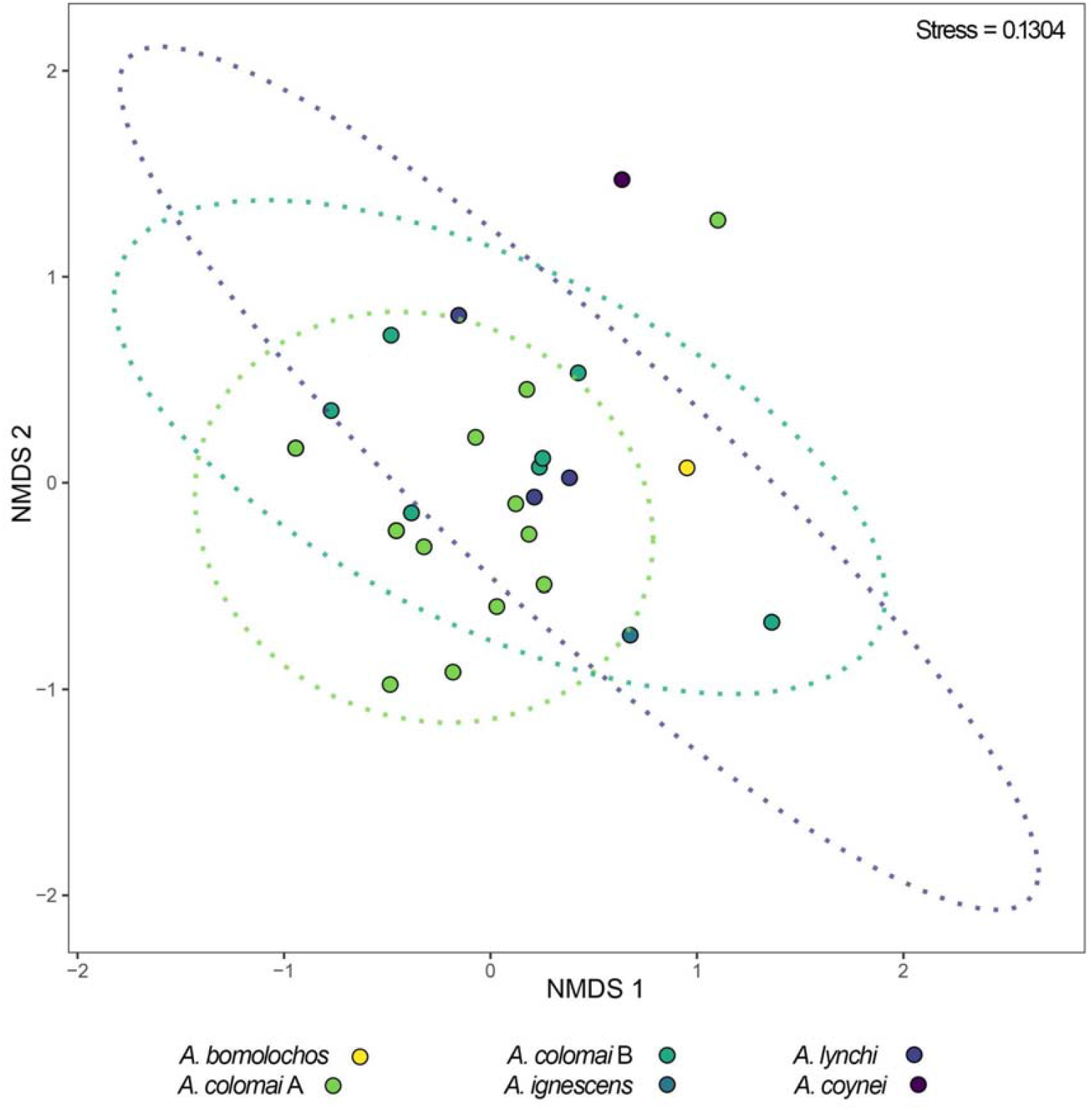
NMDS-plot of faecal metabarcoding data showing moderate interspecific differences in the dietary niche of harlequin toads but no differentiation between populations of *A. colomai* when assessed on order level. Ellipses present 0.95 % confidence area.

## Discussion

### Diet of harlequin toads

More than 100 *Atelopus* species are known, belonging to four major clades (Lötters et al. 2025). Food has been studied in only nine species, namely: *A. carbonerensis* (as *A. oxyrhynchus*) and *A. cruciger* of the Venezuelan-Andean clade; *A. ardila* (as *A. ignescens*), *A. boulengeri, A. halihelos, A. ignescens, A. sernai* and *A. varius* of the Andean-Chocó-Central American clade; and *A. planispina* (clade unknown) (Dole and Durant 1974a, Dole and Durant 1974b; Toft 1981; Gómez C. and Ramos O. 1982a, Gómez C. and Ramos O. 1982b; Valle 1984; Duellman and Lynch 1988; Ruíz-Carranza and Osorno-Muñoz 1994; González et al. 2012; Vega-Yánez et al. 2024). The diet of harlequin toads of the Sierra Nevada and the Amazonian clades remains entirely unstudied. Food items in the species mentioned was largely composed of arthropods (Hexapoda: Coleoptera, Collembola, Diptera, Homoptera, Hymenoptera; chilopods; arachnids, especially Acari; isopods) with adult ants being the most prominent prey, followed by coleopterans, both larvae and imagos. The stomachs of *A. boulengeri, A. halihelos, A. planispina* and *A. varius* almost exclusively contained ants, that of *A. carbonerensis* mainly coleopterans. Nemathelmints were frequently found in *A. ardila* (Gómez C. and Ramos O. 1982a, b), which, however, might rather represent endoparasites. Generally, high-altitude species such as *A. ardila* contained larger prey next to small items (as are most ants). This also applies to *A. carbonerensis* which consumes coleopterans of about 50% of its own body size (Dole and Durant 1974a, b). Given these data, there probably exists a tendency to feed on larger prey at higher altitudes (as *A. carbonerensis* and *A. ardila* are cloud forest and paramo species, respectively; Lötters 1996), but still in high altitude taxa high amounts of small items can be found, such as mites in the paramo species *A. ignescens* (Vega-Yánez et al. 2024; which, however, could be phoretic mites ingested with coleopterans). Dole and Durant (1974a, Dole and Durant (1974b) and González et al. (2012) reported sexual and seasonal variation in the species studied of the Venezuelan-Andean clade, while information on variation in other clades is lacking.

Our study adds the number of *Atelopus* species with dietary information to 18. While five of the species studied here belong to the Andean-Chocó-Central American clade (*A. bomolochos, A. coynei, A. ignescens, A. lynchi, A. palmatus*, with previous information only available for *A. ignescens*), *A. colomai, A. hoogmoedi, A. spumarius* and *A. tricolor* are the first species of the Amazonian clade for which digestive data are reported. In addition, microtomography of an *A. carrikeri* specimen provides the first data on diet of the basal Sierra Nevada clade. Generally speaking, the *Atelopus* food items identified by us correspond well with those of earlier studies in other species, although our analyses revealed a wider range of higher metazoan groups, for the first time reporting annelids, diplopods and onychophorans. Moreover, our dataset further favours the ‘larger prey with higher altitude’ hypothesis; however, this remains to be tested systematically once more data will become available.

### ‘Next generation’ methods for dietary studies in amphibians

Combining DNA metabarcoding and microtomography, we recovered a broad dietary niche for harlequin toads, expanding the previously known dietary spectrum of the genus. However, and despite its high resolution, a sole metabarcoding approach is currently constrained by the extreme molecular barcoding gap of tropical invertebrates, which limits the resolution of taxonomic assignment (Marques et al. 2020). Similar problems remain for ‘traditional’ morphologic examination of extracted prey items often allowing classification on higher taxonomic levels only (Vignoli and Luiselli 2012; Nielsen et al. 2017). Moreover, while read size provides a useful proxy for prey composition, it more closely reflects DNA quantity or amplification bias which is not necessarily linked to the number of individual prey items (Kennedy et al. 2020). The method is also biased towards recently ingested and large prey due to the degradation and excretion of DNA over time. Nonetheless, these limitations are equally or more pronounced in conventional dietary methods such as stomach flushing or dissection, which only reflect a very short window of predation and fail to detect soft-bodied prey that leaves few visible remains (Nielsen et al. 2017; Westeen et al. 2023). As such, metabarcoding offers a broader and less selective dietary snapshot, particularly when applied across individuals and supplemented with morphological approaches or large-scale barcoding of arthropods collected alongside the targeted predator (Nielsen et al. 2017; Pereira et al. 2021; Marques et al. 2022).

Our integrative approach highlights the power of combining metabarcoding and microtomography for reconstructing the dietary niches of threatened amphibians. While DNA metabarcoding enables standardized, non-invasive and rapid dietary profiling from contemporary faecal samples (Gillespie et al. 2023), synchrotron-based micro-CT scanning offers a non-destructive solution to assess stomach contents in preserved museum specimens, including type material. To our knowledge, this is the first study to directly visualize prey remains via µCT for dietary ecology in amphibians (Vignoli and Luiselli 2012; Nielsen et al. 2017; Ficetola et al. 2019). This novel use of microtomography complements metabarcoding by allowing non-invasive access to dietary data from historical material where molecular preservation is poor. However, imaging of chitin structures that are embedded in a complex matrix of tissues relies on advanced scanning techniques such as X-ray phase-contrast tomography or staining that reliably recover soft tissues (e.g. Metscher 2009; Sena et al. 2022).

Future methodological studies applying both techniques on the same specimens might be required to further understand methodological limitations. Emerging approaches of ancient DNA high-throughput sequencing might help to access the dietary composition of old, fluid-preserved material by employing bait-enrichment protocols from ethanol (e.g., ‘barcode fishing’ or sequence capture with metagenetic baits; cf. Scherz et al. 2020; cf. Raxworthy and Smith 2021).

### Conclusion

Our study vastly supplements the existing dietary data for near-extinct harlequin toads, offering new ecological insights critical for their conservation management. To inform ex situ management, further work will be needed, disentangling seasonal variation, sex and age-specific trophic niches as well as variation between populations. Future work will further be needed to corroborate if altitude or phylogeny shape prey spectra. The combination of faecal metabarcoding and microtomography provides a powerful and complementary framework for dietary studies, particularly suited for standardized, non-invasive sampling of threatened species and for unlocking trophic information from otherwise inaccessible museum material, including type specimens. While metabarcoding allows to capture the broad trophic niche including soft-bodied and degraded prey, microtomography provides additional quantitative information that can not be derived from metabarcoding. By providing curated and detailed code for sequence processing, our study further aims to make dietary metabarcoding more accessible for zoologists and conservationists.

## Acknowledgements

We are grateful to André Nel for identifying arthropods from imaging data. We thank Lea Groß, Felicia Hoffman and Lina Plassonke for their help with data analysis. We are grateful to Karin Fischer and Sabine Naber for their help in the lab as well as to Ambarka Giehler and Klever Velez for their support in the field. We thank Philipp Böning, Luis A. Coloma, Lara Feiler, Mark Judson, Susan Kennedy and Henrik Krehenwinkel for valuable discussions. We are grateful to Jaime Culebras for providing photographs. We thank the Ministerio del Ambiente, Agua y Transición Ecológica for issuing collection and export permits (No. MAATE-DBI-CM-2022-0261 and MAATE-ARSFC-2022-2344). AP is funded by the Research Foundation – Flanders (FWO) under a PhD Fellowship Fundamental Research (grant number 1104226N).

## Author contributions

AP, CH and SL conceptualised the study. All authors collected data. AP and CH conducted lab work. RB conducted synchrotron imaging and image processing. TH, RB and AP analysed data. AP and SL wrote the manuscript with contributions from all authors.

## Supplementary Data

All data supporting this publication is available as Supplementary Material containing sample information, additional figures and the OTU table. All custom scripts used for sequence processing and analysis are provided in Supplementary Methods. Raw sequence reads are available from Figshare under https://www.doi.org/10.6084/m9.figshare.28255364.

