## Supplementary Methods for "Metabarcoding and microtomography reveal new insights into the dietary niche of near-extinct amphibians"

1^st^ section details the Bash code for processing demultiplexed MiSeq reads

2^nd^ section details the R code for downstream filtering of OTU tables and subsequent visualization

**## Linux code for sequence processing. All characters in *italics* need to be replaced with your respective information**

\## merge all F and R files individually at once using pear:

ls | awk '{print "pear -q 20 -j 12 -v 50 -f "$0" -r "$0" -o "$0}' | grep -v _R2_ | sed 's/_R1_/_R2_/2' | sed -r 's/_S[0-9] {1,3}_L001_R1_001.fastq.gz//2' >pear.sh

chmod +x pear.sh

./pear.sh

\## quality filtering for all files at once using the fastx toolkit:

ls *assembled.fastq | awk '{print "fastq_quality_filter -Q33 -q 30 -p 90 -i "$0" | fastq_to_fasta -Q33 -o "$0}' | sed 's/assembled.fastq/fasta/2' >fast.sh

chmod +x fast.sh

./fast.sh

\## trim off primer sequences (replace all degenerate bases with a point ) and (optionally) removing short sequences. Further, all files are merged into a single .fas file:

grep -E "^*FwdPRIMERSEQUENCE.***REVERSECOMPLEMENTOFRevPRIMER*$" *fasta | sed -r 's/:[A-Z]{*FwdPRIMERLENGTH*}/\t/' | sed -r 's/[A-Z]{*RevPRIMERLENGTH*}$//' | awk 'length ($2)>=*100*{print ">"$1"\n"$2}' > *name*.fas

\## dereplicate Sequences using vsearch:

vsearch --derep_fulllength *name*.fas --output *name*.derep --sizeout

\## Cluster sequences using vsearch. With the --id option, you can set the radius (1.00 = zOTUs, 0.97 = 3% radius OTUs as in our manuscript):

vsearch --cluster_size *name*.derep --centroids *name*.OTU.fas --relabel OTU --id *0.97* --sizein --sizeout

\## then, set minsize in next step to remove rare OTUs (that are most likely derived from seq. errors):

vsearch --fastx_filter *name*.OTU.fas --minsize 5 --fastaout *name*.OTUfinal.fas

\# Creating the OTU table

vsearch --usearch_global *name*.fas --db *name*.OTUfinal.fas --id 0.97 --otutabout *name*.Tab.txt

```
## this txt can be used for further processing. It contains reads per OTU per sample file. To actually interpret our OTUs, we must now assign them to reference sequences using blastn.
## run blast search:
```

conda activate blast

blastn -db /path/nt -query name.OTUfinal.fas -num_threads 24 -max_target_seqs 10 -out name_OTUs.out -outfmt "6 qseqid sseqid pident length mismatch gapopen qstart qend sstart send evalue bitscore staxids"

conda deactivate

\## assign taxon names to OTUs:

source blast2tax_env/bin/activate

/mnt/d/Sequences/blast2taxonomy.py -i name_OTUs.out -o name_blast2tax.txt -t 6

deactivate

**## R code for filtering OTU tables and plotting**

library(dplyr)

library(ggplot2)

library(tidyverse)

library(scales)

library(openxlsx)

library(tidyr)

library(vegan)

\# diet, excl. Chordata and incl. only animals as primer pair is prone to pick up fungi.

```
TaxTab<- read.xlsx("path/Atelopus_gut_content_Taxa.xlsx", sheet = "Tabelle2") %>%
  filter(kingdom == "Metazoa") %>%
  filter(phylum != "Chordata")
```

\# Read data and merge with TaxTab

Count_OTU_diet <- read.xlsx("*path*/Atelopus_gut_content_samples.xlsx", sheet = "Tabelle2") %>%

\# Inner join with TaxTab dataframe

inner_join(TaxTab, by = "OTU_ID") %>%

\# Remove duplicated rows based on OTU_ID

distinct(OTU_ID, .keep_all = TRUE) %>%

\# Convert to dataframe

as.data.frame()

\# remove orders named Nan or NA. We filter for length smaller or equal 64 bp

\# (the expected length of amplicon after primer trimming and allow up to 30 bases lacking overlap with reference in BLAST)

\# We do this only in this step to extract the best hit from BLAST first instead of possibly only extracting a lower scored one just due to length

```
Count_OTU_diet_clean <- Count_OTU_diet %>%
  filter(sbjct_len >= 44 & sbjct_len <= 64 & perc_id >= 90000) %>%
  filter(!genus %in% c("Nan", "NA"))
```

\# Export

write.xlsx(Count_OTU_diet_clean, file = *path*/Atelopus_diet_merged.xlsx")

\# At this point I usually check the exported table visually for any potential issues,

\# manually remove OTUs that are present in negative controls and taxa that are irrelevant

\# (in our case any phyla besides Annelida, Arthropoda, Mollusca and Onychophora as other groups mostly comprise intestinal parasites or commensals not of interest for our study) and reload the table

Count_OTU_diet_clean<- read.xlsx("*path*/Atelopus_diet_merged_2025_neg_removed_forR.xlsx")

\# now to plot the data, you may summarize your replicates. The following command summarizes all samples into one column per species

merge_similar_columns <- function(data) {

\# Group columns by the first three characters of their names

grouped_cols <- unique(substr(names(data), 1, 3))

\# Iterate over grouped column names and merge them

```
  for (group in grouped_cols) {
    cols_to_merge <- grep(paste0("^", group), names(data), value = TRUE)
    if (length(cols_to_merge) > 1) {
```

\# Convert columns to numeric before summing

numeric_data <- sapply(data[cols_to_merge], as.numeric)

\# Sum values and replace the original group name

data[[group]] <- rowSums(numeric_data, na.rm = TRUE)

\# Drop the merged columns

```
      data <- data[, !names(data) %in% cols_to_merge]
    }
  }
  return(data)
}
```

\##### __________________ALTERNATIVELY______________________ #####

\# you may want to summarize replicates A & B of every individual sample and rename them by removing the _A or _B

merge_similar_columns <- function(data) {

\# Group columns by their names without the last two characters

grouped_cols <- unique(sub(".{2}$", "", names(data)))

\# Iterate over grouped column names and merge them

```
  for (group in grouped_cols) {
    cols_to_merge <- grep(paste0("^", group, ".{2}$"), names(data), value = TRUE)
    if (length(cols_to_merge) > 1) {
```

\# Sum values and replace the original group name

data[[group]] <- rowSums(data[cols_to_merge], na.rm = TRUE)

\# Drop the merged columns

```
      data <- data[, !names(data) %in% cols_to_merge]
    }
  }
  return(data)
}
```

\##### ______________________________________________________ #####

\# now apply it

Count_OTU_diet_clean <- merge_similar_columns(Count_OTU_diet_clean)

\# Convert all columns to characters, remove duplicate Taxa and calculate relative OTU abundance

```
rownames(Count_OTU_diet_clean) <- as.character(rownames(Count_OTU_diet_clean))
names(Count_OTU_diet_clean)[names(Count_OTU_diet_clean) == "Ac_"] <- "A. colomai A"
names(Count_OTU_diet_clean)[names(Count_OTU_diet_clean) == "AcF"] <- "A. colomai B"
names(Count_OTU_diet_clean)[names(Count_OTU_diet_clean) == "Ap_"] <- "A. palmatus"
names(Count_OTU_diet_clean)[names(Count_OTU_diet_clean) == "Ab_"] <- "A. bomolochos"
names(Count_OTU_diet_clean)[names(Count_OTU_diet_clean) == "Ai_"] <- "A. ignescens"
names(Count_OTU_diet_clean)[names(Count_OTU_diet_clean) == "Al_"] <- "A. lynchi"
names(Count_OTU_diet_clean)[names(Count_OTU_diet_clean) == "Aco"] <- "A. coynei"

Count_OTU_diet_class <- Count_OTU_diet_clean %>%
  mutate(across(everything(), as.character)) %>%
  select(class, starts_with("A.")) %>%
  pivot_longer(cols = -class, names_to = "Samples", values_to = "OTUs") %>%
  filter(OTUs != 0) %>%
  group_by(Samples, class) %>%
  summarize(OTUs = sum(as.numeric(OTUs))) %>%
  group_by(Samples) %>%
  mutate(Sum_OTU = OTUs / sum(OTUs)) %>%
```

\### if you want to reduce a certain taxonomic level (here: class) for simplified graphs (here: everything <= 1% abundance is merged as "Other"):

mutate(class = ifelse(Sum_OTU <= 0.01, "Other", class)) %>%

ungroup() %>%

group_by(Samples, class) %>%

summarise(n = sum(Sum_OTU)) %>%

rename(n = n) %>%

ungroup() %>%

select(class, Samples, n)

\##### ___________________________________________________ #####

\# now prior to plotting we may want to fix the order in which taxa are given by abundance. This is included in the following paragraph

\# Convert all columns to characters

```
Count_OTU_sort <- Count_OTU_diet_clean %>%
  mutate(across(everything(), as.character))

Count_OTU_sort <- Count_OTU_sort %>%
  select(class, starts_with("A.") ) %>%
  group_by(class) %>%
  summarise(across(everything(), function(x) if(is.numeric(x)) sum(x, na.rm = TRUE) else x[1]))

Count_OTU_sort <- Count_OTU_sort %>%
  select(class, starts_with("A.") ) %>%
  pivot_longer(cols = -class, names_to = "Samples", values_to = "OTUs") %>%
  filter(OTUs != 0) %>%
  #filter(OTUs !=1) %>%
  #filter(OTUs !=2) %>%
  #filter(OTUs !=3) %>%
  # filter(OTUs !=4) %>%
  group_by(Samples, class) %>%
  summarise(reads = paste(OTUs, collapse = "; ")) %>%
  ungroup() %>%
  group_by(Samples) %>%
  mutate(total_reads = sum(as.numeric(unlist(strsplit(reads, "; ")))),
         total_reads = ifelse(is.na(total_reads), 0, total_reads)) %>%
  ungroup() %>%
  mutate(Sum_OTU = as.numeric(reads) / total_reads * 100,
         Sum_OTU = ifelse(is.na(Sum_OTU), 0, Sum_OTU)) %>%
  mutate(class = ifelse(Sum_OTU <= 0.01, "zzOther", class)) %>%
  ungroup() %>%
  group_by(Samples, class) %>%
  summarise(reads = paste(reads, collapse = "; ")) %>%
  select(class, Samples, reads)
```

\# Convert 'reads' column to numeric, handling any non-numeric values appropriately

Count_OTU_sort$reads <- as.numeric(as.character(Count_OTU_sort$reads))

\# Compute the total reads for each class across all samples

species_order <- Count_OTU_sort %>%

filter(!is.na(reads)) %>% # Remove rows with NA values in 'reads' column

group_by(class) %>%

summarise(total_reads = sum(reads)) %>%

arrange(desc(total_reads)) %>%

pull(class)

\##### __________________________________________________ #####

```
plot <- Count_OTU_diet_class %>%
  ggplot() +
  aes(x = Samples, y = n, fill = factor(class, levels = species_order)) +
  geom_col(colour="black", size = 0.1) +
  labs(x = "Samples",
       y = "relative occurrence", fill = NULL) +
  scale_y_continuous(labels = scales::percent_format())+
  scale_fill_viridis_d(option = "D") +
  theme_classic() +
  coord_cartesian(expand = FALSE)+
  theme(axis.text.x = element_text(angle=45, vjust=.5, size = 12)) +
  theme(legend.position = "right",
        legend.text = element_text(size = 15),
        legend.box.margin = margin(t = 85)) +
  guides(fill = guide_legend(
    keywidth = unit(0.5, "cm"),
    keyheight = unit(0.5, "cm"),
    ncol = 1,
    #nrow = length(unique(Count_OTU_dom$family)),
    byrow = FALSE))

plot

ggsave("path/relative_abundance_class.png", plot, dpi = 500, bg = "white")
```

\##### ___________________________________________________ #####

\##### ____________________NMDS plot______________________ #####

```
df <- read.csv("path/table.txt", header = TRUE)
df <- na.omit(df)
df <- df[rowSums(df[, -1] > 0) >= 1, ]
com <- df[,2:ncol(df)]
m_com <- as.matrix(com)
set.seed(123)

nmds_result <- metaMDS(m_com, trymax = 100, distance = "bray")
#dist_matrix <- vegdist(m_com, trymax =10000, method = "jaccard")
nmds_result
plot(nmds_result)
```

##______________extract NMDS scores____________________##

nmds_scores <- scores(nmds_result, display = "sites")

nmds_data <- as.data.frame(nmds_scores)

nmds_data$Group <- df$Typ

##______________plot NMDS results with ggplot2_____________________##

```
x_limits = c(-3, 3)
y_limits = c(-3, 3)
ggplot(nmds_data, aes(x = NMDS1, y = NMDS2)) +
  geom_point(color = "black", shape = 21, size = 3, aes(fill = factor(Group))) +
  stat_ellipse(aes(color = factor(Group)), level = 0.95, linetype = 3, linewidth = 1, alpha = 0.8) +
  theme_bw() +
  theme(panel.grid = element_blank(),legend.position = "bottom") +
  labs(x = "NMDS 1", y = "NMDS 2", color = "Groups") +
  scale_fill_manual(values = c("#fde725", "#7ad151", "#22a884", "#2a788e", "#414487", "#440154")) +
  scale_color_manual(values = c("#fde725", "#7ad151", "#22a884", "#2a788e", "#414487", "#440154")) +
  coord_equal()
ggsave('path/NMDS.pdf', device = "pdf", dpi = 300)
```

##______________Calculate pairwise test for significance________________##

```
ano <- anosim(m_com, df$Typ, distance = "bray", permutations = 9999)
ano

groups <- unique(df$Typ)
pairwise_comparisons <- combn(groups, 2, simplify=FALSE)

pairwise_results <- data.frame(
  Group1  = character(),
  Group2  = character(),
  R_value = numeric(),
  p_value = numeric(),
  stringsAsFactors = FALSE
)

for (pair in pairwise_comparisons) {
  num_group1 <- sum(df$Typ == pair[1])
  num_group2 <- sum(df$Typ == pair[2])

  if (num_group1 < 2 || num_group2 < 2) {
    cat("Skipping comparison", pair[1], "vs", pair[2],
        "because one group has fewer than 2 samples.\n")
    next
  }

  indices <- df$Typ %in% pair
  subset_com <- m_com[indices, ]
  subset_typ <- df$Typ[indices]

  pair_ano <- anosim(subset_com, subset_typ,
                     distance="bray", permutations=9999)

  pairwise_results <- rbind(pairwise_results,
                            data.frame(Group1  = pair[1],
                                       Group2  = pair[2],
                                       R_value = pair_ano$statistic,
                                       p_value = pair_ano$signif))}
pairwise_results
```
